# Proinflammatory circulating extracellular vesicles from type 1 diabetes patients contribute to beta cell cytotoxicity and disease pathogenicity

**DOI:** 10.1101/2025.10.24.684420

**Authors:** Nagesha Guthalu Kondegowda, Zelda Cisneros, Daniel Roeth, Jeong-su Do, Joanna Filipowska, Rosemary Li, Shihong Zhang, Selassie Ogyaadu, Chathurani S. Jayasena, Tijana Jovanovic-Talisman, Clive Wasserfall, Yate-Ching Yuan, Helena Reijonen, Markus Kalkum, Carol J. Levy, Fouad Kandeel, Susmita Sahoo, Rupangi C. Vasavada

**Affiliations:** Arthur Riggs Diabetes and Metabolism Research Institute, City of Hope, Duarte, CA 91010, USA; Department of Medicine, Division of Endocrinology, Icahn School of Medicine at Mount Sinai, New York, NY 10029, USA; Department of Cardiology, Icahn School of Medicine at Mount Sinai, New York, NY 10029, USA; Department of Cancer Biology and Molecular Medicine, Beckman Research Institute, City of Hope, Duarte, CA 91010, USA; Diabetes Institute, College of Medicine, University of Florida, Gainesville, FL 32611, USA; Department of Computational Quantitative Medicine, City of Hope, Duarte, CA 91010, USA

**Author notes:** **Corresponding author** Rupangi C. Vasavada, Department of Translational, Research and Cellular Therapeutics Arthur Riggs Diabetes and Metabolism Research Institute, City of Hope, Room 1131E3, Gonda North; 1500 E Duarte Road, Duarte, CA 91010, USA.

**Keywords:** T1D, autoantibody-positive, islet transplant patient samples, circulating extracellular vesicles, β-cell death, NOD mice, proteomics, immune cells

## Abstract

In Type 1 diabetes (T1D), β-cell loss and autoimmunity initiate years before disease diagnosis. However, the mechanisms remain unknown. Here, we investigated the role and mechanisms of circulating extracellular vesicles (cEVs) on T1D pathogenesis and β-cell cytotoxicity.

We used cEVs isolated from healthy donors (HD), T1D, multiple autoantibody-positive (AAb+), pre- and post-islet transplantation T1D subjects, NOD-T1D mice, and T1D PBMCs. Patient T1D-cEVs significantly induced apoptosis *in vitro* in human β-cells but not α-cells compared to HD-cEVs. cEVs from AAb+ subjects and pre-diabetic mice were cytotoxic, demonstrating that cEV-induced β-cell cytotoxicity precedes T1D onset and diagnosis. cEV reduction in prediabetic NOD-T1D mice improved β-cell health implying cEV contribution to T1D development. Proteomic analysis of patient T1D-cEVs found several proinflammatory proteins, including interferon gamma, which induced β-cell cytotoxicity. Our data using cEVs from pre- and post-islet transplant and PBMC-EVs from T1D patients implicated immune cells as a potential cellular source of cytotoxic cEVs in T1D.

Collectively, our data using T1D patient samples coupled with studies in mice demonstrate that cEVs contribute to β-cell cytotoxicity and to the progression of T1D pathogenicity. Our study is one of the first to demonstrate a functional role of the proinflammatory cEV protein cargo on glucose homeostasis, β-cell health, and insulitis. Our studies on T1D-cEVs may lead to discovery of novel biomarkers and therapeutic targets for T1D.

## INTRODUCTION

Type 1 diabetes (T1D) is an autoimmune disease which results from the destruction of insulin-producing β-cells, mediated by islet autoantigen-specific T cell activation [1-4]. Emerging evidence suggests that T1D involves not only β-cell loss but also β-cell dysfunction or dedifferentiation [5-6]. T1D is characterized by four stages [7]: 1) *Precursor*, genetically predisposed but without autoimmunity; 2) *Stages 1/2*, persistent islet autoantibodies (AAbs) with asymptomatic normo- or dys-glycemia; 3) *Onset Stage 3*, with AAbs, hyperglycemia, and symptoms; and 4) *Stage 4*, established T1D. Although, islet autoimmunity and functional β-cell loss initiate years/decades before disease diagnosis in AAb+ individuals [8-9], what instigates these processes is still unknown. To date, there are no reliable predictive biomarkers that can indicate β-cell distress.

Extracellular vesicles (EVs), a family of secreted membrane-enclosed vesicles, whose molecular composition and functional characteristics reflect the pathophysiologic state of the source cell, are important mediators of inter-cellular/organ communication [10-12]. In T1D pathogenicity, there is an increased interest in tissue-derived EVs, from islets, β-cells and immune cells, their molecular cargo with a primary focus on microRNAs, and their impact on β-cell health and immune cell activation [13-15]. Circulating EVs (cEVs) in the blood have been implicated in disease pathology and pursued as predictive biomarkers for cancer, cardiovascular, neurodegenerative, and autoimmune diseases [16-20]. In T1D, cEVs induce immune activation [21], β-cell dysfunction [22], and carry specific miRNAs [22-23] distinct from healthy subject cEVs and from serum miRNAs [23]. However, the impact of T1D-cEVs on β-cell health and the mechanism by which they contribute to the pathogenesis of T1D is largely unknown. Our recent finding revealed that human T1D serum was cytotoxic to non-diabetic human β-cells [24], implying presence of cytotoxic factors in the circulation of T1D patients. Here, we investigated the role of vesicular (cEV) versus non-vesicular components in mediating humoral β-cell cytotoxicity in T1D patients.

In this study, using human samples from non-diabetic healthy donors (HD), T1D and AAb+ subjects, longitudinal samples from islet transplant T1D patients, as well as *in vivo* T1D mouse model, we reveal that cEVs contribute to T1D pathogenicity through β-cell cytotoxicity starting at the pre-diabetes stages. We show that T1D-cEVs carry several distinct proinflammatory proteins, including interferon gamma (IFNɣ), which induces β-cell cytotoxicity. We find that T1D-cEV cytotoxicity reduces in patients that have undergone islet transplantation as a therapy and are on immunosuppression, implying the immune cells as a source of cytotoxic cEVs. Indeed, EVs from PBMCs of T1D but not HD subjects induced β-cell death *in vitro*, which was mitigated in the presence of immunosuppressors, reinforcing immune cells as a potential cellular source of cytotoxic cEVs in T1D.

## RESULTS

### Human T1D-cEVs induce β-cell but not α-cell death

To identify the humoral cytotoxic components in the plasma of T1D patients, we focused our investigation on the vesicular (cEV) versus non-vesicular components. We assessed the effect of plasma and plasma-derived cEVs from T1D donors (<7 years since diagnosis) and HD best-matched for age, sex, and race/ethnicity (Table S1A) on human β-cell death. cEVs (small 30-200 nm exosome fraction) from HD and T1D donors were isolated using four distinct methods [ultracentrifugation (UC), density-based UC, modified Exoquick (removal of contaminating proteins followed by precipitation), and size exclusion chromatography (SEC)], to ensure reproducibility of the cEV data. Fractions 3-5 of the SEC method (fig. S1A) were used based on their purity and concentration. cEV size (Fig. 1A) and concentration (Fig. 1B) assessed by dynamic light scattering (DLS) or nanoparticle tracking analysis (NTA) (fig. S1B) showed no significant differences between T1D and HD subjects. cEV morphology by transmission electron microscopy (TEM) (fig. S1C) and markers by western blot analysis (fig. S1D) confirmed cEV morphology and purity from both HD and T1D donors, isolated by different methods.

**Figure 1.**
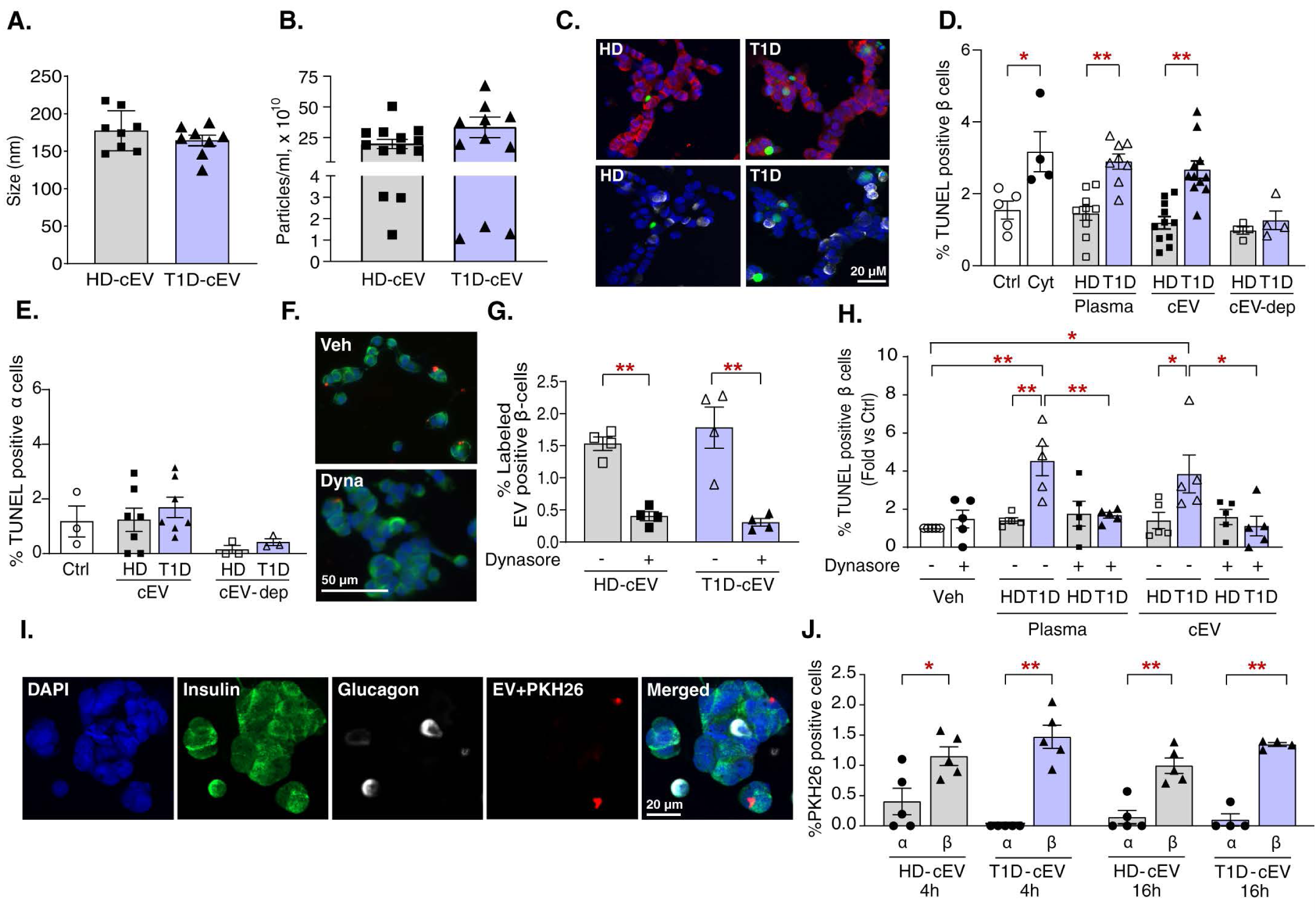
Human T1D-cEVs induce β-cell but not α-cell cytotoxicity. **(A)** Particle size (n=8/group) and **(B)** Particle concentration (n=12-13/group) of HD- and T1D-cEVs obtained using different isolation methods. **(C)** Representative confocal images of human islet cells cultured with cEVs from HD or T1D subjects and stained for insulin (red; upper panels), glucagon (white; lower panels), TUNEL (green), and DAPI (blue). **(D)** Quantification of percent TUNEL-positive β-cells in human islet cell preps (n=9) cultured in media with plasma (10%vol/vol substituting FCS) (n=7-9/group), cEVs (n=10/group) (isolated using four different methods), or EV-depleted (EV-dep) fractions (n=4/group) from HD or T1D donors. **(E)** Quantification of percent TUNEL-positive α-cells in human islet cell preps (n=3) cultured in regular media (ctrl) or with cEVs (n=7/group), or EV-dep fractions (n=3/group) from HD or T1D subjects. **(F)** Representative immunofluorescent images of human islet cells pre-cultured for 30min with vehicle (veh) or dynasore (dyna) and treated with fluorescent-labeled T1D- or HD-cEVs (red) and stained for insulin (green) and DAPI (blue). **(G)** Quantification of percent labeled HD- or T1D-cEV-positive human β-cells treated with (+) or without (-) dynasore (n=4/group). **(H)** Quantification (fold change over control) of TUNEL-positive human β-cells pre-treated with (+) or without (-) dynasore and cultured with vehicle (veh) or with T1D or HD (n=5/group) plasma or cEVs. **(I)** Representative confocal images of human islet cells treated with fluorescent-labeled T1D- or HD-cEVs (red) for 4-16h and stained for insulin (green), glucagon (white), and DAPI (blue). **(J)** Quantification of percent labeled HD- or T1D-cEV-positive human β- and α-cells at 4h and 16h post-treatment (n=4-5/group). *p<0.05, **p<0.01. White bar indicates the scale for the immunofluorescent images. The symbols in the graphs represent plasma or EV samples from individual subjects; and for the control and cytokine-treated conditions they represent distinct human islet preps. Experiments were done in duplicate; 5-10 fields and 1512±173 β-cells/sample (Fig. 1D,G,H), 1098±423 β-cells/sample (Fig. 1J), and 234±94 α-cells/sample (Fig. 1E,J) were analyzed. All data represent mean ± SEM. Statistical analysis was by Student’s t-test for comparison between two groups, and by ANOVA with Tukey’s post-hoc analysis for multiple group comparison.

To investigate T1D-cEV cytotoxicity, we isolated cEV-enriched or cEV-depleted fractions from T1D and HD plasma. Human islet cells were treated with either total plasma, cEV-enriched, or cEV-depleted fractions from both HD or T1D subjects, or with pro-inflammatory cytokine mix (as a positive control), or left untreated (ctrl) and were co-stained for insulin and TUNEL (Fig. 1C). T1D plasma-treated cells, like cytokine-treated cells, showed significantly increased β-cell death compared to HD or ctrl. Notably, T1D-cEVs (isolated using four independent methods), but not T1D-cEV-depleted, HD-cEVs or HD-cEV-depleted fractions were cytotoxic *in vitro* (Fig. 1D), implying that cEVs mediate T1D humoral cytotoxicity.

Evidence from animal models and human disease shows that β-cells and not α-cells are preferentially destroyed in T1D [25-26]. To examine the relevance of T1D-cEV cytotoxicity to the disease process, we examined the cell specificity of T1D-cEV cytotoxicity. HD- or T1D-cEV-treated human islet cell preps quantified for β-cell death (Fig. 1D) were simultaneously stained for glucagon to quantify α-cell death by glucagon-TUNEL co-staining (Fig. 1C). No effect on α-cell death was observed with T1D-cEVs versus HD-cEVs or EV-depleted control conditions (Fig. 1E). Thus, T1D-cEV cytotoxicity is specific to β-cells, suggesting relevance of cEV cytotoxicity to disease pathology.

To validate cEVs as mediators of β-cell cytotoxicity we used dynasore, an EV-uptake inhibitor that inhibits activity of dynamin, a key mediator of EV-uptake through the endocytosis pathway [27-28]. Dynasore (or vehicle)-treated human islet cells were cultured with fluorescently labeled HD- or T1D-cEVs and co-stained for insulin and DAPI (Fig. 1F). The percentage of β-cells positive for labeled HD- or T1D-cEVs was similar, and this was significantly inhibited with dynasore (Fig. 1G). We then assessed the effect of dynasore (or vehicle)-treatment on T1D plasma- and T1D cEV-induced β-cell cytotoxicity. Dynasore-treatment significantly abrogated T1D plasma or T1D-cEV-mediated β-cell cytotoxicity (Fig. 1H), implying that cEVs in the plasma mediate humoral cytotoxicity.

We next examined if differential uptake of cEVs by primary human β-versus α-cells, could explain the difference in T1D-cEV-induced cytotoxicity in these cells. Human islet cells were treated for 4h or 16h with fluorescently labeled HD- or T1D-cEVs and co-stained for insulin, glucagon and DAPI (Fig. 1I). Indeed, there were significantly higher percentages of cEV-positive β-cells versus cEV-positive α-cells at both timepoints (Fig. 1J), irrespective of the cEV source (HD or T1D). Thus, preferential cEV uptake by β-cells versus α-cells contributes to the T1D-cEV-induced β-cell-specific cytotoxicity.

### T1D-cEV carry proinflammatory cargo

To identify the molecular cargo responsible for β-cell cytotoxicity, we focused on the protein rather than the miRNA cargo, and performed a proteomic analysis, in T1D-versus HD-cEVs. A principal component analysis (PCA) showed distinct grouping of the HD-versus T1D-cEVs (Fig. 2A). The heat map indicated differential protein abundance in T1D-cEVs compared to HD-cEVs (Fig. 2B; Table S2). Pathway analysis showed upregulation of insulin secretory, proinflammatory, immune, and T1D-related pathways (Fig. 2C), with IFN-γ being the cytokine with the highest significant log-fold change (Fig. 2D; Table S2) in T1D-versus HD-cEVs. ELISA analysis confirmed a significant increase in IFN-γ levels in T1D plasma and cEVs versus HD (Fig. 2E). To test the function of IFN-γ in T1D-cEVs, we treated human islet cells with HD- or T1D-cEVs and co-stained for insulin, DAPI, and phSTAT1, a key transcription factor, activated by IFN-γ (Fig. 2F). The percentage of phSTAT1-positive β-cells was significantly higher with T1D-cEV treated cells compared to HD-cEV-treated cells or controls (Fig. 2G). To determine if the IFN-γ in T1D-cEVs contributes to their cytotoxicity, we pre-treated human islet cells with the IFN-γ inhibitor, emapalumab (EMP) [29] prior to cEV treatment. EMP significantly reduced T1D-cEV-induced β-cell death, (Fig. 2H), implying IFN-γ as a mediator of T1D-cEV cytotoxicity.

**Figure 2.**
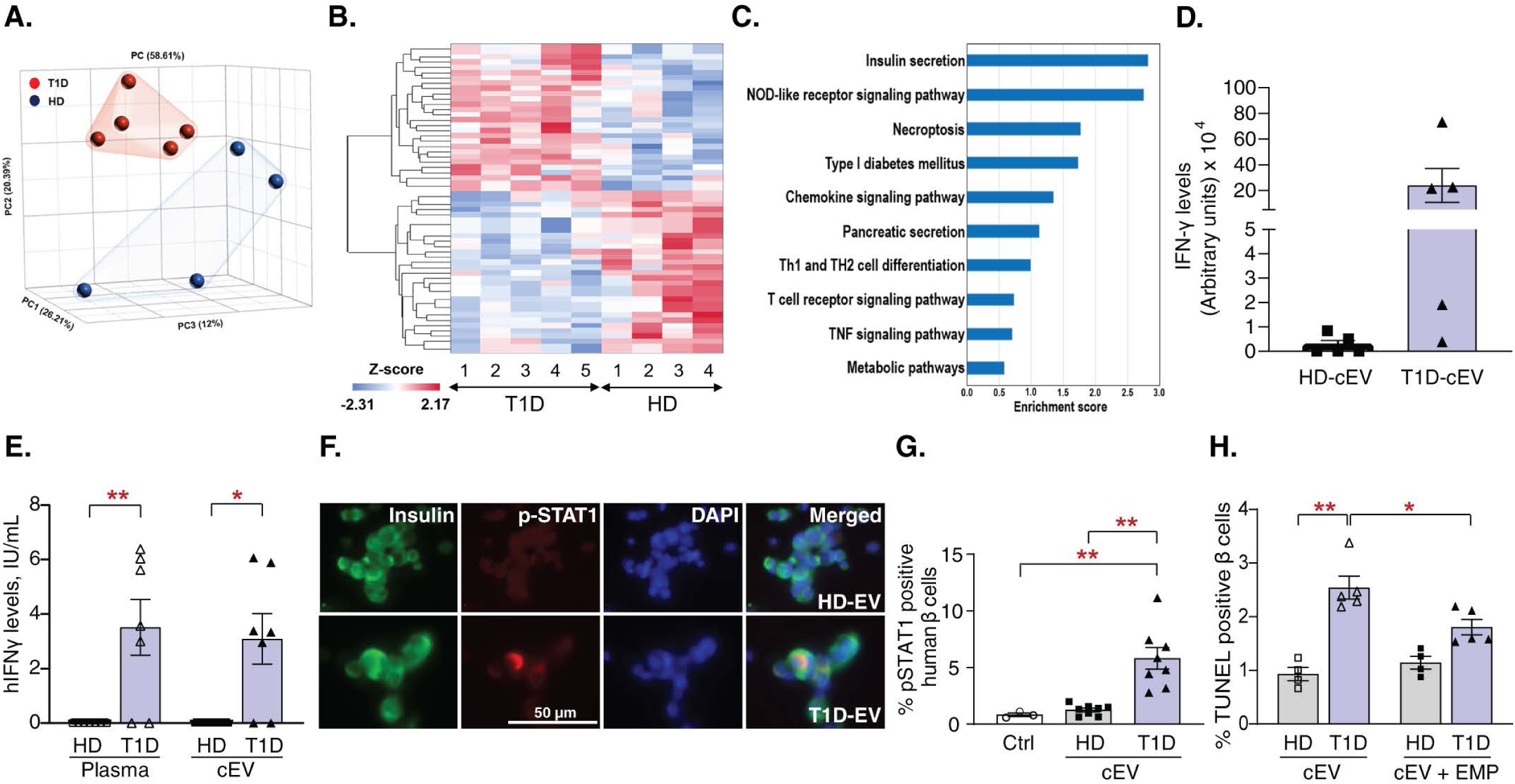
Human T1D-cEVs carry proinflammatory cargo, including IFN-γ, which mediates β-cell cytotoxicity. Proteomic analysis of cEVs from T1D (n=5) and HD (n=4) subjects showing **(A)** Principal component analysis (PCA), **(B)** Heatmap of the differential cargo, **(C)** Pathway analysis, and **(D)** IFN-γ levels. **(E)** IFN-γ levels measured by ELISA in plasma and cEVs of HD and T1D donors (n=7/group). **(F)** Representative images of human islet cells treated with cEVs from HD or T1D subjects and stained for insulin (green), phSTAT1 (red), and DAPI (blue). **(G)** Quantification of percent phSTAT1-positive β-cells in human islet cell preps (n=2) treated with HD- or T1D-cEVs (n=8/group), or in untreated control cells. **(H)** Quantification of percent TUNEL-positive β-cells in human islet cell preps pre-treated with the IFN-γ inhibitor, EMP or vehicle (veh), and subsequently treated with HD- (n=4) or T1D-cEVs (n=5). *p<0.05, **p<0.01. White bar indicates the scale for the immunofluorescent images. Individual symbols in the graphs represent samples from individual subjects, or individual human islet preps for the control conditions. Experiments were done in duplicate; 5-10 fields and 1814±75 β-cells/sample were analyzed for Fig. 2G,H. All data represent mean ± SEM. Statistical analysis was by Student’s t-test for comparison between two groups, and by ANOVA with Tukey’s post-hoc analysis for multiple group comparison.

### cEV reduction improves glucose homeostasis in NOD mice

To evaluate the *in vivo* contribution of cEVs to T1D pathogenesis, we used female NOD mice, a widely used T1D murine model that spontaneously develops autoimmune diabetes [30-31]. We first verified if NOD mouse cEVs, like human T1D donor samples (Fig. 1D), induced β-cell death *in vitro*. We treated C57Bl6 mouse islet cell cultures with cEVs isolated from the sera of female NOD mice that remained non-diabetic, had early onset diabetes, or had established diabetes and assessed β-cell death by insulin-TUNEL co-staining. Consistent with human data (Fig. 1D), cEVs from early onset and established diabetic NOD mice showed elevated β-cell death compared to non-diabetic mouse cEVs (Fig. 3A), establishing the NOD mouse as a relevant model to assess the contribution of cEVs in T1D pathogenesis *in vivo*. We used GW4869, a neutral sphingomyelinase 2 inhibitor, the most widely used pharmacological agent to block EV biogenesis and reduce EV release [32]. Six-week-old prediabetic female NOD mice were injected with vehicle or GW4869 (5 µg/g body weight, based on a dose-response study) every alternate day for six weeks, with body weight and blood glucose measured weekly, intraperitoneal glucose tolerance test (IPGTT) performed at D38 of treatment, blood sampled at D0 pre-treatment and at D42 end of treatment, and pancreata harvested at the end of the study (fig. S2A). The cEV concentration was significantly reduced with GW4869 versus vehicle treatment at D42, or versus D0 pre-treated mice (Fig. 3B), indicating that GW4869-treatment reduced cEV levels. Body weight was unaltered during treatment (fig. S2B). The GW4869-treated group showed significantly lower blood glucose by mixed model analysis during the treatment compared to the vehicle-treated group (Fig. 3C). IPGTT showed significantly improved glucose tolerance in the GW4869- versus vehicle-treated group (Fig. 3D), and the area under the curve (AUC) between the two groups was almost significant (p=0.058) (fig. S2C). The insulin response to the IPGTT was poor in the vehicle-treated NOD mice (Fig. 3E), as expected [33], however, plasma insulin at the 15-minute time point in the GW4869-treated mice was significantly increased relative to the 0-minute time point in vehicle-treated mice (Fig. 3E). Plasma insulin at D42 was not statistically different between the two groups (fig. S2D). Reducing cEV levels in pre-diabetic NOD female mice improved glucose homeostasis.

**Figure 3.**
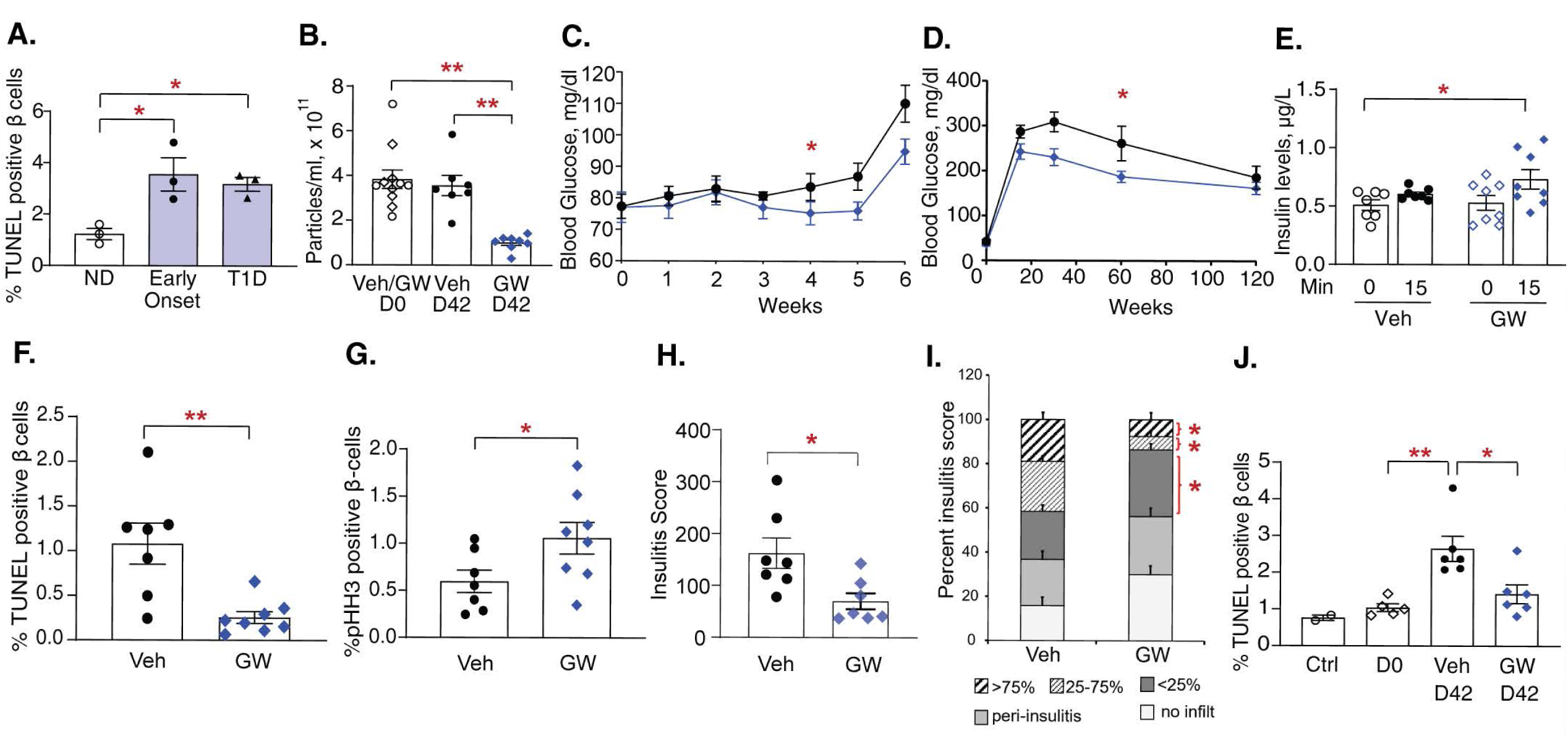
Reducing cEV levels in pre-diabetic NOD mice improves glucose homeostasis, β-cell health, and insulitis. **(A)** Percent TUNEL-positive β-cells in mouse islet cells treated with cEVs (n=3/group) isolated from the sera of female NOD mice that remain non-diabetic (ND), or have early onset diabetes, or have established diabetes (T1D). **(B)** Concentration of cEV particles from the sera of mice at pre-treatment (D0) or at the end of treatment (D42) from the vehicle (Veh)- and GW4869 (GW)-treated groups. Vehicle- (black symbols and line) and GW4869-treated (blue symbols and line) mice (n=7-8/group) were assessed for **(C)** Blood glucose, **(D)** IPGTT, and **(E)** Plasma insulin at 0- and 15-min time-points during the IPGTT. Pancreatic sections from vehicle and GW4869-treated mice were quantified for **(F)** percent TUNEL-positive β cells; **(G)** percent pHH3-positive β cells; **(H)** overall insulitis score; and **(I)** percent insulitis in specific quartiles. **(J)** Percent TUNEL-positive β-cells in mouse islet cells cultured in media with FCS (ctrl) or media in which the FCS was substituted with serum (10% vol/vol) from vehicle- or GW4869-treated NOD mice at D0 or D42 of treatment (n=5-6 mice/condition). *p<0.05, **p<0.01. The symbols in the graphs represent samples from individual mice; and for the control they represent mouse islet preps. Experiments were done in duplicate; 7-19 islets and 1966±235 β-cells/sample were analyzed. All data represent mean ± SEM. Statistical analysis was by Student’s t-test for comparison between two groups, by mixed model analysis for Fig. 3, C and D, and by ANOVA with Tukey’s post-hoc analysis for multiple group comparison.

### cEVs negatively impact NOD mouse β-cells *in vivo*

Based on the β-cell cytotoxic effects observed with rodent (Fig. 3A) and human (Fig. 1D) T1D-cEVs, we hypothesized that reducing cEV levels with GW4869-treatment in NOD mice will improve β-cell health *in vivo*. TUNEL-insulin co-staining in pancreatic sections (fig. S2E) showed a significant decrease in β-cell death in the GW4869- versus vehicle-treated mice at D42 (Fig. 3F). Assessment of β-cell replication by insulin-phospho-histone 3 (pHH3) co-staining (fig. S2F) showed a significant increase in β-cell proliferation in the GW4869- versus vehicle-treated mice (Fig. 3G). Insulitis, scored on H&E-stained sections (fig. S2G), showed a significant decrease in the overall insulitis score [34] (Fig. 3H), which was reflected in a significant reduction in the higher quartiles of insulitis (>25%) and a significant increase in the lower quartiles of insulitis (<25%) in GW4869- versus vehicle-treated mice (Fig. 3I). To further validate that NOD mouse cEVs induce β-cell cytotoxic effects, we tested the effect of serum from vehicle- and GW4869-treated mice on β-cell cytotoxicity *ex vivo*. Interestingly, there was a significant increase in β-cell death when cells were cultured in serum from pre-diabetic D42-vehicle-treated versus from D0-pre-treated mice (Fig. 3J). However, serum from D42-GW4869-treated mice, with reduced cEV levels (Fig. 3B), showed significantly reduced β-cell death versus D42-vehicle-treated mice (Fig. 3J), demonstrating that cEVs induce β-cell death and contribute to T1D pathology.

### cEV cytotoxicity precedes T1D diagnosis

The loss of functional β-cell mass and immune cell activation initiates well before clinical T1D diagnosis [35]. Our finding that cEV-induced β-cell cytotoxicity exists in *pre-diabetic* NOD mice (Fig. 3, F and J), suggested that cEV phenotype is altered prior to clinical T1D diagnosis. To test this in humans, we obtained blood from AAb+ donors (≥2 AAb+) (Table S1B) at high risk for T1D [36-37], from two independent sources. Human islets cells were co-cultured with plasma, cytokines (positive control) or cEVs and assessed for β-cell death. Although plasma from AAb+ donors induced β-cell death it was not statistically different from HD serum (Fig. 4A). In contrast, cEVs isolated from AAb+ donors significantly increased β-cell death versus HD-cEVs (Fig. 4A). Taken together, this implies that cEVs acquire a β-cell cytotoxic phenotype prior to clinical disease onset.

**Figure 4.**
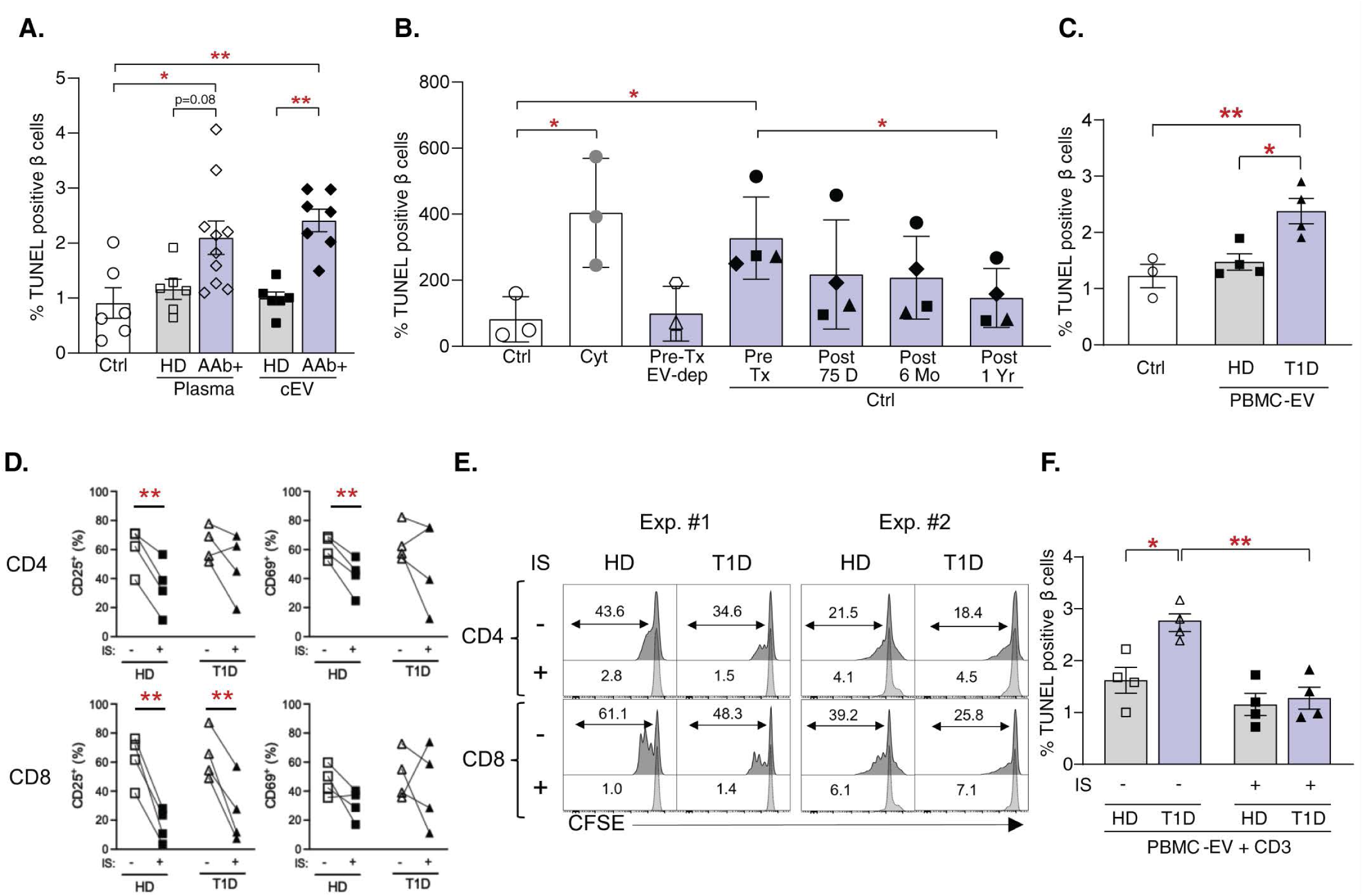
β-cell cytotoxicity mediated by cEVs and PBMC-derived EVs in varied clinical settings of T1D. Quantification of percent TUNEL-positive β-cells in human islet preps (n=3-6) cultured in media with FCS (Ctrl), **(A)** or media in which the FCS was substituted with serum/plasma (10% vol/vol), or treated with cEVs isolated from a subset of the samples of HD and multiple AAb+ subjects (n=6-10); **(B)** or treated with cEVs or cEV-depleted (EV-dep) fractions isolated from serum samples of T1D (n=4) patients before islet transplantation (Tx) (Pre-Tx) and 75 days, 6 months, and 1 year post-islet Tx, or with cytokine mix (Cyt) as a positive control for cell death; **(C)** or treated with EVs isolated from PBMCs of HD and T1D subjects (N=4/group). PBMCs from HD and T1D donors were treated with anti-CD3 ± immunosuppressants (IS) and assessed for **(D)** percent immune cell activation markers CD25 and CD69 (N=4/group), and **(E)** percent proliferating cells (indicated by the number) (N=2/group) in CD4 (upper panel) and CD8 (lower panel) cells. **(F)** Percent β-cell death in human islet preps treated with EVs isolated from PBMC culture (+anti- CD3 ± IS) of HD and T1D subjects. *p<0.05, **p<0.01. The symbols in the graphs represent samples from individual subjects; and for the control and cytokine-treated conditions they represent distinct human islet preps. Experiments were done in duplicate; 5-15 fields and 1523±248 β cells/sample were analyzed. All data represent mean ± SEM. Statistical analysis was by Student’s t-test for comparison between two groups, and by ANOVA with Tukey’s post-hoc analysis for multiple group comparison.

### cEV-cytotoxicity reduces post-islet transplantation

A clinically relevant question is whether cEV cytotoxicity changes in T1D patients that have undergone islet transplantation as a therapy and have been exposed to immunosuppressants. We isolated cEVs at four distinct time-points (pre-transplant; 75 days, 6 months, and 1-year, post-islet transplant), from four T1D patients who underwent islet cell transplantation (Table S1C). Human islet cells showed a significant increase in β-cell death when treated with T1D-cEVs versus EV-depleted fraction from pre-transplant patients or compared to untreated controls (Fig. 4B). Thus, pre-transplant T1D-cEVs phenocopied cEVs from T1D donors (Fig. 1D). Remarkably, the β-cell cytotoxic effect of cEVs from the same patients gradually declined with increasing time post-islet transplantation, with a significant reduction in β-cell death observed with cEVs at 1-year post-islet transplantation versus cEVs pre-islet transplantation (Fig. 4B). This implies that cEV characteristics indeed change after islet transplantation. The major changes in T1D patients, post-islet transplantation, are improved blood glucose regulation through enhanced insulin secretion from functional β-cells, and continuous suppression of the immune system. One explanation for the blunted cEV β-cell cytotoxicity is that immune cells are a source of cytotoxic cEVs, and thus, immunosuppression after islet transplantation reduces the production of cytotoxic cEVs.

### PBMCs, source of cytotoxic cEVs in T1D

To address whether immune cells may be a source of cytotoxic cEVs in T1D, we isolated PBMC-derived EVs from T1D and HD donors (Table S1D) and tested their cytotoxic effect on human β-cells. T1D-PBMC-EVs significantly increased β-cell death versus HD-PBMC-EVs (Fig. 4C), indicating that like cEVs, immune cell-derived EVs from T1D donors also induce β-cell cytotoxicity. To determine if immunosuppression of activated PBMCs could reduce their EV β-cell cytotoxicity, cultured PBMCs from T1D and HD donors were activated with anti-CD3 antibody in the presence or absence of immunosuppressants. Tacrolimus and sirolimus, immunosuppressants used in T1D patients undergoing islet transplantation [38], were used *in vitro,* at concentrations found in the circulation of post-islet transplant T1D recipients. Effectiveness of the immunosuppressants on activated T1D and HD PBMCs was demonstrated using flow cytometry showing a decrease in immune cell activation markers (Fig. 4D) and a reduction in proliferation (Fig. 4E) in CD4+ and CD8+ T-cells. EVs from activated T1D PBMCs but not HD PBMCs significantly induced β-cell death (Fig. 4F). Importantly, immunosuppressant-treatment significantly blunted the β-cell cytotoxic effect of T1D-PBMC-EVs to basal levels (Fig. 4F). Our data shows that T1D-PBMC-EVs, derived basally or after immune cell activation, are cytotoxic to β-cells, unlike HD-PBMC-EVs. Cytotoxicity of T1D-PBMC-EVs is significantly reduced upon immunosuppression. Using clinical samples from T1D patients, our data implies PBMCs as a potential source of cytotoxic cEVs in T1D.

## DISCUSSION

Despite recent advances in our understanding of the pathogenesis of T1D, we still cannot identify the contributors to the initiation or progression of the disease. The interplay between genetic susceptibility, environment, β-cell health, and activation of the autoimmune response, plays a key role in T1D. Therefore, cellular signaling and crosstalk between tissues through circulation is critical for disease manifestation. EVs have gained prominence in this regard, through their regulation of normal physiology and pathophysiology, such as cancer, cardiomyopathy, and neurodegenerative diseases [16-20]. However, the contribution of cEVs to the pathogenesis of T1D is not known. In T1D, the composition of serum/plasma [39-40] and PBMC characteristics [41-42] are significantly changed compared to healthy control subjects, implying that humoral components could participate in the disease process. Our recent study shows that serum from T1D, but not control subjects, is cytotoxic to human β-cells [24], implying that cEVs in T1D could play a detrimental role on β-cell health, contributing to disease pathology.

To test this, we used cEVs from T1D subjects (<7 years since disease diagnosis) and non-diabetic HD subjects, best matched for age, sex, and race/ethnicity, as well as mouse cEVs from non-diabetic and T1D mice. To ensure robustness of our findings, human cEVs were isolated using four distinct methods that vary in the purity and abundance of EV output, with the SEC and density-gradient UC methods giving the highest purity but low abundance of EVs, versus the modified Exoquick [43] and UC methods giving higher abundance but lower purity EVs [44]. With all four methods, the larger EV fraction was first separated out, and the smaller (exosome) EV fraction was used for the studies. We demonstrate that T1D but not control cEVs of human and mouse origin are cytotoxic to β-cells *in vitro*. Furthermore, we show that the humoral cytotoxicity of plasma from T1D patients is mediated through cEVs. We find that cEVs specifically kill human β-but not α-cells, mirroring the cellular specificity seen in disease pathogenesis [25-26]. One possible explanation for the β-cell specific cytotoxicity of T1D-cEVs is that β-cells are more susceptible to stress and cell death relative to α-cells based on their gene expression profile [26] and lower levels of antioxidants [45]. Another possible explanation could be that EVs are preferentially taken up by β-versus α-cells, which has not been examined thus far. Indeed, our studies show that there is a preferential uptake of both HD- and T1D-cEVs by human β-versus α-cells, which could in part explain the β-cell specific cytotoxic effects of cEVs. The mechanism of EV uptake by β-cells, whether it is specific receptor mediated uptake, or through endocytosis, or by plasma membrane fusion, is not known, and will be examined in future studies.

To understand the mechanism behind the T1D-cEV-induced β-cell cytotoxicity, we examined the molecular cargo of cEVs. Unlike other studies that have assessed miRNAs in cEVs, this is the first proteomic analysis on HD- and T1D-cEVs. We identified distinct protein cargo in T1D-cEVs enriched in proinflammatory and immunomodulatory pathways, including T1D, chemokine, and secretory pathways amongst others. Pro-inflammatory IFNɣ was the cytokine with the highest significant log-fold change in T1D-cEVs, and this increase was corroborated independently by an ELISA in T1D plasma and cEVs. IFNɣ has proinflammatory and cytotoxic effects on the β-cell, regulates CD8+ T cell homing to the islet, can amplify islet autoimmunity, and control proliferation of autoreactive T cells [46-47]. The IFNɣ in T1D-cEVs was functional and contributed to the cEV-induced cytotoxicity, as demonstrated through activation of the downstream phSTAT1 pathway, and reduction of β-cell death with the IFNɣ inhibitor, in human β cells. How IFNɣ, a cytosolic protein, that should typically reside in the lumen of T1D cEVs, mediates its effect on the target β-cell is not known. The mechanism could be through binding of IFNɣ to the T1D-cEV corona [48], or through IFNɣ binding to its receptor present on the EV surface, as observed in EVs from neuronal stem cells [49], or through its re-release as internalized cargo from the recipient cell [50]. Future studies will determine the mechanism by which IFNɣ in the cEVs induces β-cell death as well as identify other mediators relevant in T1D-cEV-induced cytotoxicity.

Although our *in vitro* studies on T1D patient-derived cEVs underscore the clinical relevance of our findings, they do not directly address the contribution of cEVs to the disease process. To investigate that, we turned to a mouse model of spontaneous T1D, the NOD/Ltj female mice. We first confirmed that this T1D mouse is relevant to our hypothesis, using *in vitro* studies, that phenocopied our observations with the T1D patient-derived cEVs. To understand the role of cEVs in T1D pathogenesis *in vivo*, we treated *prediabetic* 6-week-old female NOD/Ltj mice for six weeks with vehicle or GW4869, a sphingomyelinase inhibitor that impairs EV biogenesis and release. Although this is not a specific EV inhibitor it is a widely used tool, both *in vitro* and *in vivo* studies, in the EV field [32]. The GW4869 regimen chosen (5µg/g every alternate day) was based on our pilot study (not shown) using different doses and frequency of administration. The GW4869-treatment reduced cEV levels by ∼75% at the end of the treatment period. GW4869-treatment resulted in a significant reduction in blood glucose, an improvement in glucose tolerance, with no significant changes in plasma insulin, a significant reduction in β-cell death, an increase in β-cell proliferation, and a reduction in the level of insulitis compared to the vehicle-treated controls. Our *ex vivo* studies using serum from these mice corroborated our *in vivo* observations of cEV-induced β-cell cytotoxicity. The reduction in insulitis could be a result of the reduced β-cell death observed in these mice, as β-cell apoptosis creates a microenvironment of inflammation that induces immune cell infiltration and activation [51], or a direct effect of cEVs on immune cell activation and function. Future studies will test whether GW4869 treatment will have a greater impact on older mice that are more likely to develop diabetes, whether GW4869-treatment can reverse diabetes in mice, and whether cEVs directly impact immune cell activation.

The finding that reducing cEV levels in *prediabetic* NOD mice improves glucose and β-cell homeostasis implies that cEVs contribute to the disease process even before the onset of T1D, and thus, are relevant to the development and progression of the disease. To examine this concept in humans, we assessed the β-cell cytotoxic effects of plasma and cEVs from AAb+ (≥2 AAbs) donors with no clinical disease diagnosis of T1D. AAb+ cEVs, but not AAb+ plasma nor HD-cEVs, significantly increased β-cell death. This implies that cEVs acquire a proinflammatory phenotype prior to clinical disease onset, and hence, are likely to contribute to the underlying ongoing disease progression.

To examine whether the cEV phenotype is altered in T1D patients that have undergone islet transplantation as a therapy and are under an immunosuppressive regimen, we used longitudinal samples of cEVs pre- and post-islet transplantation from T1D patients. There was a significant reduction in the cytotoxicity of cEVs isolated one-year post-islet transplantation compared to the cEVs from pre-islet transplantation, indicating that the cEVs had undergone a change in their phenotype. This could be due to the presence of functional β-cells, improved glucose control, and/or the immunosuppressive regimen used post-islet transplantation. We hypothesized that one possible cellular source of the cytotoxic cEVs in T1D could be the immune cells, and therefore, a reduction in immune cell activation through immunosuppression could lead to reduced cEV cytotoxicity. We tested this using T1D and HD PBMC-derived EVs and found that only T1D-PBMC-EVs but not HD-PBMC-EVs, were cytotoxic to β-cells, and this cytotoxicity was significantly reduced with immunosuppression, further validating the likelihood that PBMCs are a significant source of cytotoxic cEVs in T1D.

The strengths and innovation of this study include the use of cEVs from patient samples, including people with T1D, multiple AAb+ subjects, and longitudinal samples from T1D patients that have undergone islet transplantation. Using four distinct EV isolation methods resulting in the same functional phenotype lends credence and weight to our findings. Our *in vitro* studies demonstrate a critical role of cEVs in T1D disease pathology, through their selective cytotoxicity to human β-versus α-cells, mediated at least partially through their selective uptake by β-cells. Our *in vivo* findings in the NOD mouse define a causative role of cEVs in T1D pathogenesis. The first proteomic analysis of T1D-cEVs identify proinflammatory cargo in patient-derived T1D-cEVs and its causative effect on β-cell cytotoxicity provides novel mechanistic insight. Our studies in mouse and AAb+ individuals demonstrate that cytotoxic cEVs appear prior to the clinical diagnosis of disease, indicating an involvement in disease development and progression. The abating of the T1D-cEV cytotoxicity in T1D patients after islet transplantation indicates that the cEV phenotype can change based on the metabolic, immune, and/or treatment profile of the patient. Finally, identifying PBMCs as a potential source of the cytotoxic cEVs in T1D patients can pave the way towards future novel therapeutics.

The limitations of the study include the non-specificity of action of the EV uptake and biogenesis inhibitors, dynasore and GW4869, respectively. Although these inhibitors have been used extensively in EV research, they also impact other biological processes, and hence, the results should be interpreted with caution. Future studies using different inhibitors with distinct modes of action can further validate these findings. Outcomes from the current study highlight many interesting and important questions. For example, what is the mechanism of the β-cell specific cytotoxic effect of cEVs? How early in the disease process do cEVs acquire the cytotoxic phenotype? Can the molecular cargo in cEVs serve as biomarkers for early disease detection? Do cEVs impact other cells and tissues such as immune cells, and other peripheral target organs? What specific cell type(s) in PBMCs is the source of cEVs? Could reducing cEV levels delay disease progression in animal models or in people? Future studies will address these and other relevant questions.

## METHODS

See Supplemental Material for detailed methods.

## FUNDING

This work was supported by National Institutes of Health grants R01DK125856 (R.C.V. and S.S.) and R01DK133885 (R.C.V. and H.R.), Juvenile Diabetes Research Foundation (JDRF)- no. 1- INO-2017-438-A-N (R.C.V. and S.S.); Wanek Family Project to Cure Type 1 Diabetes at City of Hope (R.C.V.); AR-DMRI Pilot Study (R.C.V. and H.R.).

## CREDIT AUTHORSHIP CONTRIBUTION STATEMENT

*NGK*: Conceptualization, Data curation, Formal analysis, Investigation, Methodology, Project administration, Visualization, Writing - original draft; and Writing - review & editing; *ZC*: Data curation, Investigation, Methodology, Writing - review & editing; Figure layout and editing; *DR*: Data curation, Investigation, Software, Writing - review & editing; *JD*: Data curation, Investigation, Writing - review & editing; *JF*: Data curation, Writing - review & editing; *RL*: Data curation, Writing - review & editing; *SZ*: Data curation, Writing - review & editing; *SO*: Data curation, Sample acquisition; *CSJ*: Sample acquisition, Writing - review & editing; *TJT*: Methodology, Writing - review & editing; *CW*: Sample acquisition, Writing - review & editing; *YCY*: Data curation, Investigation, Software, Writing - review & editing; *HR*: Sample Acquisition, Supervision, Funding acquisition, Writing - review & editing; *MK*: Data curation, Supervision, Software, Writing - review & editing; *CJL*: Sample acquisition, Supervision, Writing - review & editing; *FK*: Conceptualization, Supervision, Sample acquisition, Writing - review & editing; *SS*: Conceptualization, Supervision, Methodology, Funding acquisition, Writing - review & editing; *RCV*: Conceptualization; Resources, Formal analysis, Supervision, Validation, Project administration, Writing - original draft, Writing - review & editing, and Funding acquisition.

## ACKNOWLEDGMENTS

We are grateful to the donors of human islets and blood samples; to the NIDDK-supported Integrated Islet Distribution Program, Southern California Islet Resource Center and Prodo Laboratories for providing human islets; to Dr. Mark Atkinson (University of Florida) for providing human serum/plasma samples (NIH grant AI42288); to Dr. Navendu Vasavada for expertise on statistical analysis (http://astatsa.com/); to the microscopy core (Dr. Brian Armstrong), and pathology core supported by the National Cancer Institute of the National Institutes of Health under grant number P30CA033572, at COH for assistance and use of their facilities; and to COH Translational Bioinformatics resources, support, and training for data exploration, visualization, analysis, and discovery.

## DECLARATION OF COMPETING INTEREST

The authors declare no competing interests.

## DATA AVAILABILITY

Data will be made available on request.

## SUPPLEMENTARY MATERIAL

### Supplementary Methods

#### Human plasma, serum, and PBMCs

Blood was drawn from human donors after receiving informed consent in accordance with institutional guidelines based on the approved Institutional Review Board (IRB) protocols. Whole blood was processed to obtain plasma (from purple-topped EDTA or blue-topped sodium citrate tubes), serum, and PBMCs as per standard protocols. Samples were stored at -80°C or under liquid nitrogen as appropriate number and volume of aliquots to avoid repeated freeze and thaw cycles. Samples from T1D (<7 years since disease diagnosis) and healthy donor (HD) matched for sex, age, and race/ethnicity for plasma/serum (Table S1A) and PBMCs (Table S1D) were obtained at City of Hope, CA, and Icahn School of Medicine at Mount Sinai, NY (IRB #20477, #18156, and #15-01137), from multiple (>2AAbs) AAb+ subjects (Table S1B) were obtained at University of Florida, FL, and Pacific Northwest Research Institute (PNRI), WA, from T1D patients who have undergone islet transplantation (Table S1C) were collected under City of Hope IRB-approved Protocol #12446 (ClinicalTrials.gov ID: NCT01909245). Leftover frozen EDTA plasma, at four different time points (pre-transplant, 75 days, 6 months, and 1-year post-transplant), were used for retrospective analysis.

#### EV isolation

cEV isolation from plasma or serum was performed using four different methods, ultracentrifugation (UC), sucrose gradient UC, modified Exoquick [43] and size exclusion chromatography (SEC). EV isolation from PBMCs was performed using overnight UC. For all methods, centrifugation steps were performed at 4°C, and the samples were centrifuged at 500g for 10 min and then at 2000g for 10 min to remove cell debris and apoptotic bodies. Larger vesicles were removed by centrifuging at 10,000 rpm for 30 min and the remaining supernatant was used for EV isolation. For UC, 1 ml plasma diluted with PBS at a ratio of 2:1 (PBS to plasma volume) was pre-clarified and then ultracentrifuged at 100,000 × g for 90 min to pellet the EVs. The EV pellet was resuspended in PBS, washed by a second ultracentrifugation at 100,000 × g for 90 min, and finally resuspended in 120 µL of PBS. The samples were stored at −80°C until further use. Sucrose gradient EV isolation was performed under sterile conditions using a sucrose/D₂O gradient ultracentrifugation protocol. First, cell debris and larger vesicles were removed as mentioned above. The resulting supernatant was carefully layered onto a sucrose/D₂O solution and ultracentrifuged at 100,000 x g for 90 min. EVs were concentrated at the interface of the gradient layer. This EV-enriched fraction was carefully collected and washed with PBS, followed by a final ultracentrifugation step at 100,000 x for 90 min to pellet the EVs. The final EV pellet was resuspended in 100 µL of PBS and stored at -80°C until further use. For the modified ExoQuick method pre-clarified plasma (100-200 µl) was mixed with an equal volume of thromboplastin D and incubated for 15 min at 37°C. Supernatants were collected following centrifugation at 13,000 × g for 10 min at 4°C, mixed with 50-100 µl of ExoQuick reagent (EXOQ5A, System Biosciences, Pal Alto, CA) (Table S3) and incubated overnight at 4°C. The preparation was spun down at 1,500 × g for 25 min at 4°C. The top portion of the supernatant (∼50 ul) was carefully collected and considered as EV-depleted fraction, and the rest of the supernatant was discarded. The tube with pellet was recentrifuged at 1500 x g, 4 ^◦^C, for 5 min to remove the leftover supernatant. EV-enriched pellet was resuspended in 50-100 µl of PBS. SEC was performed with 500 µl of pre-clarified plasma using the column (IZON Science, qEVoriginal columns, 70 nm, Medford, MA) and automatic fraction collector as per the manufacturer’s instructions. The 500 µl fractions (#3-5) containing EVs were stored at -80°C.

#### EV characterization

Particle number and size distribution of the EV preparations were tested by NTA using a NanoSight NS300 system (Malvern Technologies, Malvern, UK) configured with a 488 nm laser and a high sensitivity scientific CMOS camera. Samples were diluted (1:500) in particle-free PBS to an acceptable concentration, according to the manufacturer’s recommendations. Samples were analyzed under constant flow conditions with a flow rate of 30 at 25°C and 3×60 s videos were captured with a camera level of 12. Data was analyzed using NTA 3.1.54 software with a detection threshold of 5. DLS measurements of EV preparation were performed with Brookhaven 90Plus Particle Size Analyzer (Brookhaven Instruments Corporation). 6 µl sample was diluted in PBS to a total volume of 50 µl, and 6 measurement runs (1 min each) were performed with standard settings (Refractive Index = 1.331, viscosity = 0.89, temperature = 25°C). Only measurements with an average count rate in the 300-500 kcps range were further analyzed. Data acquisition and processing were performed using Brookhaven Particle Sizing Software v3.79 (Brookhaven Instruments Corporation) and displayed as a number size distribution. For structural analysis, isolated EVs were deposited onto formvar/carbon-coated copper EM grids (10 µl on each grid) for 20min. The vesicle-coated grids were washed three times with PBS and then fixed with 2.5% glutaraldehyde for 10min. After washing with 3 drops of distilled water, the grids were stained with 2% uranyl acetate for 15 min and air dried for 20 min. Transmission electron microscopy (TEM) was performed using a FEI Tecnai 120 microscope operating at an accelerating voltage of 120 keV. EV markers such as Alix, Flotillin 1, tetraspanin CD9, (Table S3) were analyzed by western blot analysis.

#### EV labeling

EV labeling was done using PKH26 dye as per the manufacturer’s instructions (Sigma-Aldrich, #PKH26GL-1KT, St. Louis, MO). Briefly, EV pellet was diluted in 1 ml of diluent C from the kit, 2 µl of pre-spun (16,000 g for 5 min) PKH26 dye was added and mixed gently with continuous pipetting for 30 sec., allowed to stand for 5 min at room temperature, and then 3 ml of PBS was added. This was loaded onto 1 ml of D_2_O/sucrose solution, ultracentrifuged at 40,400 rpm for 90 min, supernatant discarded, EV pellet washed s in 4 ml of PBS, ultracentrifuged at 40,400 rpm for 90 min, and re-suspended in 50 µl of PBS.

#### Mouse and human islet isolation and cell culture

Mouse islets (from C57Bl6/J mice) were isolated by collagenase digestion and histopaque gradient separation and cultured in complete medium [RPMI supplemented with 5.5mM glucose, 10% FCS, and 1% penicillin-streptomycin (pen/strep)], as described previously [24]. Human islets (Table S4) were obtained from the Integrated Islet Distribution Program (IIDP), Prodo Labs, and Southern California Islet Resource Center. Rodent and human islets were cultured in complete media at least overnight before they were hand-picked under the microscope for treatment and analysis. Islet cell cultures were prepared by 10-min trypsinization with intermittent pipetting. Cells from 50 islet equivalents (IEQs; 1 IEQ=125 μm diameter) were plated on coverslips in a 24-well plate initially in a small volume (50 μl) of media for 2h to allow attachment, after which 1 ml of media was added [24; S1].

#### Cellular assays

For the EV uptake analysis, human islet cell cultures were treated for 4 or 16h with PKH26 labeled EVs (6.4 x 10^7^ particles/ml) or PKH26 dye alone under dark conditions and then fixed with 4% paraformaldehyde and stored at 4°C before insulin/glucagon/DAPI co-staining. For assessing the effect of serum, plasma and EVs on β- or α-cell cytotoxicity, mouse or human islet cells were cultured for 24h in media containing FCS (control), or in media where the FCS was substituted with either serum or plasma (10% v/v), or in media containing EV-free FCS and treated with mouse or human EVs (5-20 µl of EVs or 50-100 µg of EV protein/ml), or with EV-depleted fraction, or treated with a mix of human cytokines (IL1β 50U, 0.714 ng/ml; IFNγ 1000U, 50 ng/ml; and TNFα 1000U, 13.15 ng/ml) (R&D Systems, Minneapolis, MN) as positive control for cell death. For EV uptake inhibition human islet cell cultures were pre-treated with vehicle or dynasore (Selleckchem, #S8047, Houston, TX) 50 µM for 30 min, before culturing with plasma or cEVs for 24h. To test for ph-STAT1 activation human islet cell cultures were treated with cEVs for 24h, fixed and stained for ph-STAT1/insulin/DAPI. To examine the effect of IFNγ, human islet cells were pre-treated with vehicle or the IFNγ inhibitor emapalumab (EMP) (Selleckchem, #A2041, Houston, TX) (Table S3) (100 µg/ml) for 4h, before treating with cEVs for 24h. For hPBMC-derived EV treatment, similar particle numbers (∼4-8 x 10^8^ particles/ml) were used to treat human islet cells for 24h. Cells were fixed in 4% paraformaldehyde for 20 min and co-stained for TUNEL, insulin, glucagon, and 4’,6-diamidino-2-phenylindole (DAPI).

#### Immunofluorescence staining in mouse and human islets *in vitro*

β- and α-cell death in human or mouse islet cells was assessed by staining for insulin (1:1000, DAKO, Sata Clara, CA; 1:1000, Abcam, Waltham, MA), glucagon (1:500, Abcam, Waltham, MA), TUNEL (Promega, Indianapolis, IN) and DAPI (Life Technologies), using Alexa Fluor 594 Goat anti-Guinea Pig or Alexa Fluor 594 Goat anti-Rabbit, and Alexa Fluor 647 Goat anti-Mouse (Life Technologies, Carlsbad, CA) as secondary antibodies (Table S3), and quantified as percentage of TUNEL-insulin-positive to total insulin-positive or TUNEL-glucagon-positive to total glucagon-positive cells [24; S1]. For EV uptake assays human islets were stained as indicated above for insulin, glucagon and DAPI, and the percentage of PKH26 labeled EV (red)-insulin-positive to total insulin-positive or PKH26 labeled EV (red)-glucagon-positive to total glucagon-positive cells were quantified. For ph-STAT1 immunostaining human islet cells were stained for ph-STAT1 (Ser727) (1:100, Cell Signaling Technology, Danvers, MA), insulin, and DAPI, and the percentage of ph-STAT1-insulin-positive to total insulin-positive cells were quantified.

#### Proteomic Analysis

Proteomic cargo analysis was done on cEVs from T1D and HD subjects (N=4-5 each). All solvents, reagents and chemicals used for LC/MS analysis were of Mass Spectrometry-grade quality. 30 μg of EV proteins were treated with 10% SDS in 100 mM triethylammonium bicarbonate (TEAB, Thermo Fisher Scientific) at a ratio of 1:1 and subsequently reduced for 30 min at 37°C with 5 mM (2-carboxyethyl) phosphine (TCEP) and alkylated at room temperature with 30 mM iodoacetamide for 45 min. Next, proteins were precipitated using suspension traps (S-Trap micro, Protifi) and sample cleanup was performed according to the manufacturer’s instructions. Digestion was performed in the S-Traps for 16h with 3 μg Trypsin/LysC (Promega) per sample, at 37°C. Peptides were reconstituted in 0.1% formic acid (FA). Mass spectrometry was performed on an orbitrap Exploris 480 instrument (Thermo Fisher) equipped with an Easy-nLC 1200 HPLC system, a 75 µm × 50 cm PepMap RSLC C18 analytical column (2 μm particle size, 100Å pore size), and an Easy-Spray ion source. Peptide nano-LC separation was performed on a 3h gradient from 100 % buffer A (0.1% FA in water) to 68% buffer A and 32% buffer B (80% acetonitrile, 0.1% FA in water) at a flow rate of 300 nL/min. Spectra were acquired using data-independent acquisition (DIA) using m/z 4 overlapping staggered windows at a width of m/z 8, in a precursor range from m/z 400 to 1000. Precursor scans were performed at a resolution of 120,000 and DIA MS/MS scans were performed with a resolution of 30,000, an AGC target of 1000% and a normalized collision energy of 27%. Thermo raw files were deconvoluted and converted to mzml format using MsConvert (v3.0.21101). Library-free searches were performed using DIA-NN (v1.8.1). Statistical analysis was performed using MS stats (v 4.0). Proteomic data analysis was performed by using Illumina Partek™ Flow™ (Version 12.7.0) (Illumina, San Diego, USA) which facilitates pre- and post-QA/QC, Principal Component Analysis (PCA), normalization using rank-based quantile method and added one before log2 transformation using raw protein measurement, differential protein expression using ANOVA, differential protein expression comparing T1D vs HD control was filtered by a significance threshold of |fold change|≥ 1.5; p-value ≤ 0.05 for downstream volcano plot visualization, heatmap, and KEGG/GSEA pathway analysis among 9 human samples, excluding one HD sample indicated as an outlier from PCA analysis.

#### Protein Assays

Human IFNγ in plasma and cEVs of T1D and HD subjects and mouse insulin from serum of NOD mice were assayed using ELISA (Enzyme Linked Immunosorbent Assay) kits for human IFNγ (KAC1231 Invitrogen, Frederick, MD) and mouse insulin (Mercodia, Inc, Winston Salem, NC) [24; S1], respectively. For western blot analysis, protein extracts (20-40μg) from cEVs and EV-depleted fractions were analyzed by immunoblotting using antibodies against Alix, Flotillin 1, and tetraspanin CD9 (Table S3). Western blot analysis was performed by treating the protein blotted PVDF membrane with appropriately diluted primary antibodies overnight individually and then treated with HRP-conjugated species-specific secondary antibodies for one hour at room temperature. Membrane was probed by treating with chemiluminescent agent to visualize the protein signal as bands.

#### Animal studies

For *in vitro* studies, islets were isolated from 4-6-month-old male C57BL/6 mice (Jackson Laboratories, Bar Harbor, ME). cEVs were isolated from the sera of female NOD/ShiLtJ mice (Jackson Laboratories, Bar Harbor, ME) that did not develop diabetes (defined as blood glucose ≥250mg/dl for 2 consecutive days), or that had early onset T1D (blood glucose ≥250mg/dl for 3 consecutive days), or established T1D (blood glucose ≥250mg/dl for 30 days). For *in vivo* studies, pre-diabetic 6-week-old female NOD/ShiLtJ mice were injected intraperitoneally (ip) with vehicle (saline) or 5.0 µg/g body weight of GW4869 (Cayman Chemical, Ann Arbor, MI), every alternate day for 42 days. Body weight and blood glucose were measured every week and IPGTT was performed at the end of the study. Blood samples were collected through facial vein before (D0) and at the end (D42) of treatment. At termination of the study, pancreata were harvested to examine changes in endogenous β-cell death, proliferation and insulitis. Animal studies were performed with the approval of, and in accordance with, guidelines established by Beckman Research Institute, City of Hope, Duarte, CA, and principles of laboratory animal care were followed.

#### Glucose homeostasis

Blood glucose was measured weekly on tail snips using a portable glucometer (Freestyle Lite, Amazon), and IPGTT was performed in the last week of treatment. Before performing IPGTT, mice were fasted for 16-18h and injected with 2 g glucose/kg body weight. Blood glucose was measured pre- and at regular intervals starting at 15 to 120 mins post-glucose treatment. Plasma insulin was measured on blood drawn at fasting, after 15 min of glucose challenge during IPGTT, and before termination of the study [24].

#### β-cell death, proliferation, and insulitis in mice

Pancreatic sections were stained with antibodies against insulin (1:1000, Dako, Carpenteria, CA) and either TUNEL (Promega, Indianapolis, IN) to measure β-cell death, or pHH3 (1:500, Millipore, Billerica, MA) to measure β-cell proliferation, after antigen retrieval with high pressure cooker steam in citrate buffer for 20 min, using an immunofluorescence secondary antibody. β-cell death was quantified as percentage of TUNEL-insulin to total insulin-positive cells and β-cell proliferation was quantified as percentage of pHH3-insulin to total insulin-positive cells [24]. Changes in insulitis levels in pancreata was analyzed in H&E-stained pancreatic sections, by manually scoring the degree of insulitis [34] for every islet in a blinded fashion and assessing at least 24.71±2.70 islets/mouse from multiple sections.

#### Human PBMC culture and assays

Frozen PBMCs from HD and T1D donors were thawed slowly in the water bath at 37°C, and media with 50% serum was slowly added while gently mixing the thawed cells. RPMI complete media was added similarly and centrifuged at 200g for 5 minutes. Cells were washed 2 times with 5% serum in RPMI. PBMC pellet was resuspended in RPMI complete media and the cells were allowed to recover for at least 3h at 37°C before seeding in the 48-well cell culture plate. PBMCs were either left untreated for 72h and culture media collected for EV isolation, or activated with anti-CD3 (OKT3, Ultra-LEAF™ Purified anti-human CD3 Antibody, Bio legend, 1 µg/ml) and treated with either vehicle or immunosuppressants (IS) (Sirolimus, Selleckchem, 6 ng/ml; Tacrolimus, Selleckchem, 6 ng/ml) for 72h and cells as well as culture media collected for functional characterization and EV isolation, respectively. The media was centrifuged at 500 g for 10 min to remove the cell debris and stored at -80°C until EV preparation.

Immune activation markers and proliferation was assessed in CD4 and CD8 T cells of PBMC. PBMCs were harvested on day 3 after activation and stained with surface-binding antibodies against CD4-PE, CD8-Brilliant Violet 785, CD25-Brilliant Violet 421, CD69-PE/Cy7, (BioLegend, San Diego, CA) (Table S3). Fc receptor blocking was performed using Human TruStain FcX (BioLegend). Staining was carried out for 15 minutes at 4°C. Flow cytometry analyses included exclusion of doublets. Fluorescence intensities were acquired using the NovoCyte Quanteon flow cytometer (Agilent Technologies) and analyzed with FlowJo software (Tree Star, Ashland, OR). For proliferation assay, PBMCs were labeled with carboxyfluorescein succinimidyl ester (CFSE, Thermofisher Scientific), and FACS analyzed.

#### Statistical analysis

All *in vitro* experiments were done in duplicate, and all analyses was performed in a blinded fashion. Data are expressed as means ± SEM. Statistical significance was considered at p<0.05 and p<0.01, for single and double symbols, respectively, determined by unpaired two-tailed Student’s t-test for comparison between two groups; by paired two-tailed Student’s t-test for comparison between PBMC treated and untreated groups; by One-way Analysis of Variance (ANOVA) with Tukey’s post-hoc HSD (http://astatsa.com/) for comparison between more than two groups; by mixed model analysis for repeated measures for the glucose measurements over time; and using MS stats (v 4.0) for the proteomic analysis.

## Supplementary Figure Legends

**Supplementary Figure 1.**
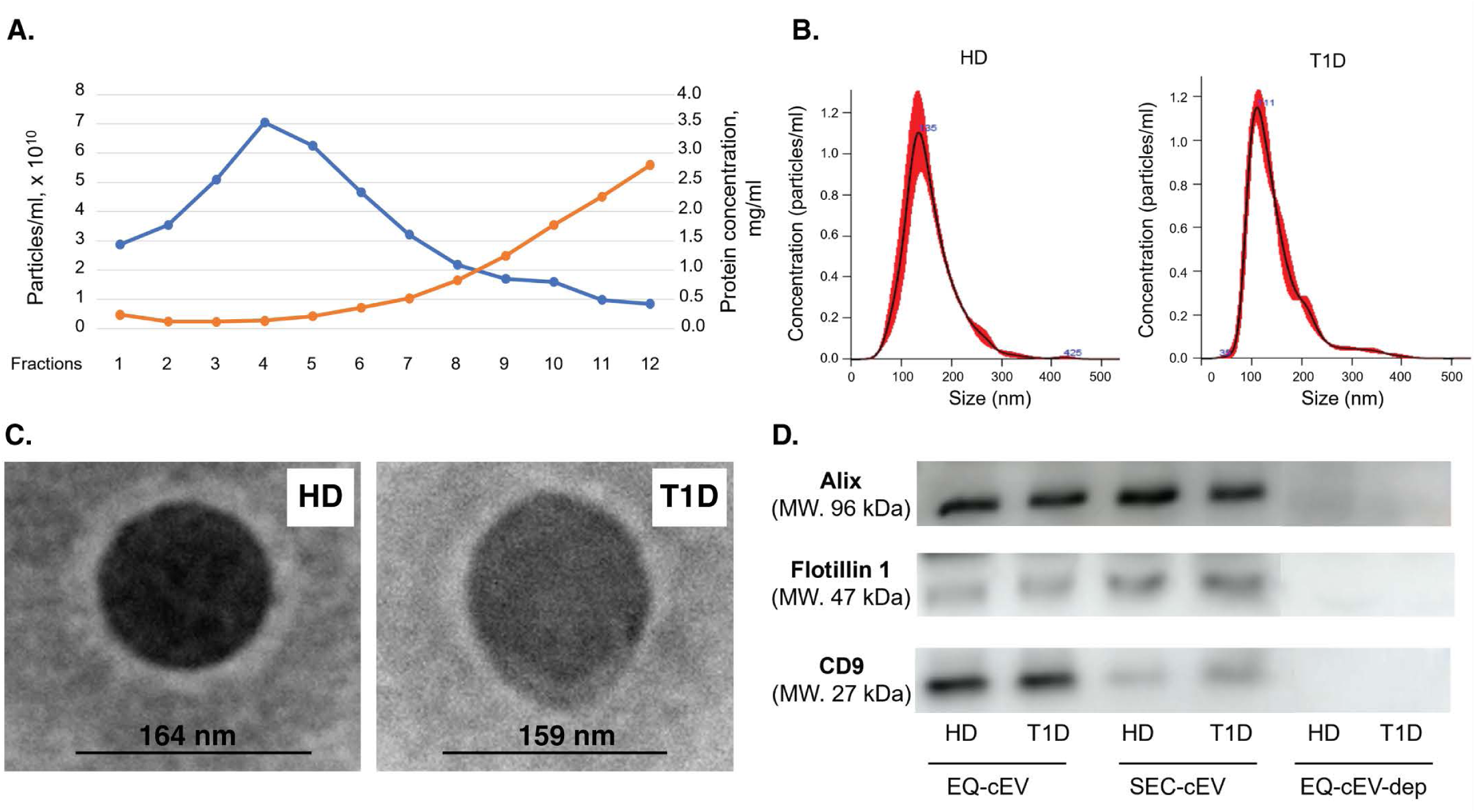
Characterization of T1D- and HD-cEVs. **(A)** Representative graph indicating particle concentration (blue line) and protein contamination (brown line) of plasma-derived cEV prep by the SEC method. **(B)** Representative graph of NTA analysis indicating size and concentration of particles from HD- and T1D-cEV preps. **(C)** Representative TEM image of HD- and T1-cEV preps. **(D)** Representative western blot analysis of EV markers, flotillin, Alix and CD9, in HD- and T1D-cEVs and cEV-depleted fractions.

**Supplementary Figure 2.**
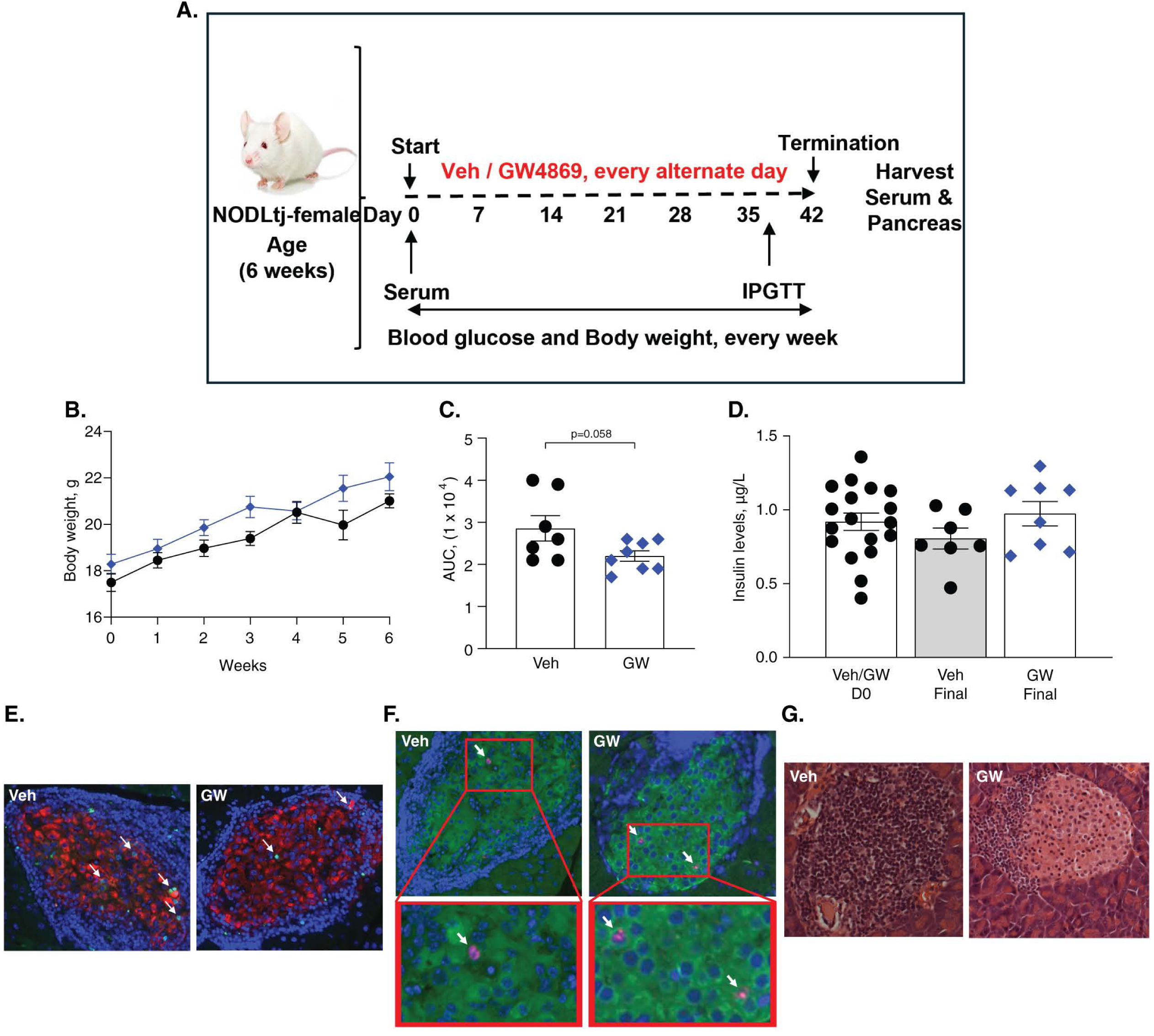
Glucose and β-cell homeostasis in GW4869-treated pre-diabetic NOD mice. **(A)** Schematic representation of the experimental design for the treatment of six-week-old prediabetic female NOD mice with vehicle (n=7) or GW4869 (5 µg/g body weight) (n=8) every alternate day for six weeks, assessing body weight and blood glucose weekly, IPGTT at D38, blood samples at D0 and D42, and pancreata at the end of the study (D42). Vehicle- and GW4869-treated mice (n=7-8/group) were assessed for **(B)** Body weight, **(C)** IPGTT AUC, and **(D)** Plasma insulin at D0 (pre-treatment) and D42 (end of treatment). Representative image of pancreatic sections from vehicle- and GW4869-treated mice **(E)** stained for insulin (red), TUNEL (green), and DAPI (blue); **(F)** stained for insulin (green), pHH3 (red), and DAPI (blue), with the boxed insets magnified in the panels below; and **(G)** stained with H&E. *p<0.05, **p<0.01. Individual symbols in the graphs represent data from individual mice. All data represent mean ± SEM. Statistical analysis was by Student’s t-test for comparison between two groups, and by ANOVA with Tukey’s post-hoc analysis for multiple group comparison.

**Supplementary Table 1A:** Characteristics of HD and T1D subjects.

| **Deidentified Subject ID** | **Sex** | **Age at Enrollment** | **Ethnicity** | **Years with Diabetes** |
| --- | --- | --- | --- | --- |
| **HD – Subjects** |  |  |  |  |
| 1 | M | 22 | Caucasian, Non-Hispanic | NA |
| 2 | M | 23 | African American, Non-Hispanic | NA |
| 3 | M | 23 | Caucasian, Non-Hispanic | NA |
| 4 | M | 25 | Caucasian, Non-Hispanic | NA |
| 5 | M | 29 | Caucasian, Non-Hispanic | NA |
| 6 | M | 31 | Caucasian, Non-Hispanic | NA |
| 7 | M | 54 | Caucasian, Hispanic | NA |
| 8 | M | 57 | Caucasian, Non-Hispanic | NA |
| 9 | F | 29 | Caucasian, Non-Hispanic | NA |
| 10 | F | 37 | Caucasian, Non-Hispanic | NA |
| 11 | F | 45 | More than one race, Hispanic | NA |
| 12 | F | 57 | Asian | NA |
| **T1D – Subjects** |  |  |  |  |
| 13 | M | 18 | Caucasian, Non-Hispanic | <1 |
| 14 | M | 22 | Caucasian, Non-Hispanic | 1 |
| 15 | M | 23 | Caucasian, Non-Hispanic | 1 |
| 16 | M | 24 | African American, Non-Hispanic | <1 |
| 17 | M | 28 | Caucasian, Non-Hispanic | <1 |
| 18 | M | 28 | Caucasian, Non-Hispanic | 2 |
| 19 | M | 28 | Caucasian, Non-Hispanic | <1 |
| 20 | M | 33 | Caucasian, Non-Hispanic | 3 |
| 21 | M | 51 | Caucasian, Hispanic | <1 |
| 22 | M | 59 | Caucasian, Non-Hispanic | 1 |
| 23 | F | 30 | Caucasian, Hispanic | 2 |
| 24 | F | 38 | Caucasian, Hispanic | <1 |
| 25 | F | 53 | Caucasian, Non-Hispanic | 3 |
| 26 | F | 56 | Caucasian, Non-Hispanic | 1 |
| 27 | F | 68 | Caucasian, Non-Hispanic | 2 |

**Supplementary Table 1B:** Characteristics of AAb+ and HD subjects.

| **Deidentified**<br>**Subject ID** | **Sex** | **Age at Enrollment** |
| --- | --- | --- |
| **Aab+ Subjects** |  |  |
| AB-1 | M | 10 |
| AB-2 | M | 10 |
| AB-3 | M | 17 |
| AB-4 | F | 34 |
| AB-5 | F | 43 |
| AB-6 | M | 11 |
| AB-7 | M | 16 |
| AB-8 | M | 17 |
| AB-9 | F | 10 |
| AB-10 | F | 13 |
| AB-11 | F | 14 |
| AB-12 | F | 17 |
| **HD Subjects** |  |  |
| HD-1 | F | Unknown |
| HD-2 | Unknown | Unknown |
| HD-3 | M | Unknown |
| HD-4 | F | Unknown |
| HD-5 | Unknown | Unknown |
| HD-6 | Unknown | Unknown |

**Supplementary Table 1C:** Characteristics of T1D subjects who have undergone islet transplant.

| **Subject ID** | **Sex** | **Ethnicity** | **Age at transplantation** | **Duration of T1D** |
| --- | --- | --- | --- | --- |
| Tx-1 | M | Caucasian, Non-Hispanic | 20 | 15 |
| Tx-2 | M | Caucasian, Non-Hispanic | 67 | 41 |
| Tx-3 | F | Caucasian, Non-Hispanic | 33 | 24 |
| Tx-4 | F | Caucasian, Non-Hispanic | 52 | 25 |

**Supplementary Table 1D:** Characteristics of HD and T1D PBMC donors.

| **Deidentified Subject ID** | **Sex** | **Age at Enrolment** | **Ethnicity** | **Years with Diabetes** |
| --- | --- | --- | --- | --- |
| **HD PBMC Donor Subjects** |  |  |  |  |
| PB-1 | M | 22 | Caucasian, Non-Hispanic | N/A |
| PB-2 | M | 36 | Caucasian, Non-Hispanic | N/A |
| PB-3 | F | 25 | Caucasian, Non-Hispanic | N/A |
| PB-4 | F | 30 | African American, Non-Hispanic | N/A |
| **T1D PBMC Donor Subjects** |  |  |  |  |
| PB-5 | M | 28 | Caucasian, Non-Hispanic | 2 |
| PB-6 | M | 34 | Caucasian, Non-Hispanic | <1 |
| PB-7 | F | 24 | African American, Non-Hispanic | 3 |
| PB-8 | F | 29 | Caucasian, Non-Hispanic | 2 |

**Supplementary Table 2:** Differential protein cargo T1D- vs HD-cEVs.

| **No.** | **Human Protein Name** | **UniProtKB ID** | **Log2 (Ratio)**<br>**(T1D vs Control)** | **P-value**<br>**(T1D vs Control)** |
| --- | --- | --- | --- | --- |
| 1 | CBX1 | P83916 | 12.794 | 0.0098 |
| 2 | IFNG | P01579 | 6.031 | 0.0252 |
| 3 | TEBP | Q15185 | 4.714 | 0.0030 |
| 4 | VWF | P04275 | 1.967 | 0.0277 |
| 5 | CAPR1 | Q14444 | 1.770 | 0.0283 |
| 6 | VAT1L | Q9HCJ6 | 1.640 | 0.0423 |
| 7 | APMAP | Q9HDC9 | 1.638 | 0.0159 |
| 8 | DEF1 | P59665 | 1.616 | 0.0073 |
| 9 | DEF3 | P59666 | 1.616 | 0.0073 |
| 10 | KV116 | P04430 | 1.546 | 0.0407 |
| 11 | NCAM1 | P13591 | 1.395 | 0.0417 |
| 12 | A4 | P05067 | 1.342 | 0.0067 |
| 13 | SERB1 | Q8NC51 | 1.324 | 0.0456 |
| 14 | FA5 | P12259 | 1.272 | 0.0003 |
| 15 | PI42B | P78356 | 1.243 | 0.0304 |
| 16 | PTPRZ | P23471 | 1.221 | 0.0269 |
| 17 | LCAT | P04180 | 1.171 | 0.0471 |
| 18 | SETD7 | Q8WTS6 | 1.099 | 0.0386 |
| 19 | SNP25 | P60880 | 1.014 | 0.0133 |
| 20 | TRFL | P02788 | 1.006 | 0.0082 |
| 21 | KTNB1 | Q9H079 | 0.916 | 0.0088 |
| 22 | AT1A3 | P13637 | 0.901 | 0.0196 |
| 23 | PF4V | P10720 | 0.875 | 0.0274 |
| 24 | SHBG | P04278 | 0.835 | 0.0354 |
| 25 | RHOB | Q3ZBW5 | 0.711 | 0.0130 |
| 26 | SAA2 | P0DJI9 | 0.667 | 0.0273 |
| 27 | SAA4 | P35542 | 0.667 | 0.0273 |
| 28 | PSB4 | P28070 | 0.627 | 0.0493 |
| 29 | CLUS | P10909 | 0.572 | 0.0148 |
| 30 | GNAO | P09471 | 0.571 | 0.0407 |
| 31 | SNAB | Q9H115 | 0.550 | 0.0369 |
| 32 | PROC | P04070 | 0.547 | 0.0448 |
| 33 | PSA5 | P28066 | 0.502 | 0.0164 |
| 34 | SEPT7 | Q16181 | 0.404 | 0.0472 |
| 35 | PSA7 | O14818 | 0.395 | 0.0287 |
| 36 | PSMA8 | Q8TAA3 | 0.395 | 0.0287 |
| 37 | RALA | P11233 | 0.379 | 0.0363 |
| 38 | RALB | P11234 | 0.379 | 0.0363 |
| 39 | RAB3A | P20336 | 0.372 | 0.0464 |
| 40 | CETP | P11597 | 0.347 | 0.0434 |
| 41 | RAB1B | Q9H0U4 | 0.345 | 0.0363 |
| 42 | CO8A | P07357 | 0.334 | 0.0224 |
| 43 | GBB2 | P62879 | 0.296 | 0.0279 |
| 44 | OLA1 | Q9NTK5 | -0.305 | 0.0399 |
| 45 | UGPA | Q16851 | -0.309 | 0.0408 |
| 46 | C1RL | Q9NZP8 | -0.416 | 0.0042 |
| 47 | UBC12 | P61081 | -0.429 | 0.0024 |
| 48 | UBE2N | P61088 | -0.449 | 0.0301 |
| 49 | UE2NL | Q5JXB2 | -0.449 | 0.0301 |
| 50 | VATG2 | O95670 | -0.464 | 0.0379 |
| 51 | BLMH | Q13867 | -0.470 | 0.0120 |
| 52 | LDHA | P00338 | -0.502 | 0.0163 |
| 53 | COR1A | P31146 | -0.506 | 0.0064 |
| 54 | ARP5L | Q9BPX5 | -0.526 | 0.0136 |
| 55 | PAPS1 | O43252 | -0.546 | 0.0365 |
| 56 | KV401 | P06312 | -0.549 | 0.0489 |
| 57 | 6PGD | P52209 | -0.565 | 0.0159 |
| 58 | PA1B3 | Q15102 | -0.629 | 0.0478 |
| 59 | A0A0C4DH41-HV461 | A0A0C4DH41 | -0.646 | 0.0316 |
| 60 | ERP29 | P57759 | -0.658 | 0.0358 |
| 61 | IF5A1 | P63241 | -0.728 | 0.0416 |
| 62 | IF5AL | Q6IS14 | -0.728 | 0.0416 |
| 63 | OSBL1 | Q9BXW6 | -0.732 | 0.0322 |
| 64 | LIS1 | P43034 | -0.749 | 0.0412 |
| 65 | UB2D2 | P62837 | -0.752 | 0.0380 |
| 66 | UBE3D | Q7Z6J8 | -0.752 | 0.0380 |
| 67 | STIP1 | P31948 | -0.752 | 0.0252 |
| 68 | PSPC1 | Q8WXF1 | -0.769 | 0.0228 |
| 69 | PTPRS | Q13332 | -0.769 | 0.0212 |
| 70 | MAOM | P23368 | -0.825 | 0.0054 |
| 71 | KV133 | P01594 | -0.841 | 0.0287 |
| 72 | CH10 | P61604 | -0.864 | 0.0222 |
| 73 | S4R3N1 | S4R3N1 | -0.864 | 0.0222 |
| 74 | HARS1 | P12081 | -0.913 | 0.0344 |
| 75 | SYHM | P49590 | -0.913 | 0.0344 |
| 76 | BAIP2 | Q9UQB8 | -0.921 | 0.0059 |
| 77 | DNM1L | O00429 | -0.944 | 0.0291 |
| 78 | FKBP3 | Q00688 | -1.244 | 0.0441 |
| 79 | HV343 | A0A0B4J1X8 | -1.371 | 0.0156 |
| 80 | P0DSN7 | KVD37 | -1.399 | 0.0107 |
| 81 | ARRB1 | P49407 | -1.428 | 0.0034 |
| 82 | MOES | P26038 | -1.546 | 0.0292 |
| 83 | YBOX1 | P67809 | -1.554 | 0.0331 |
| 84 | YBOX2 | Q9Y2T7 | -1.554 | 0.0331 |
| 85 | YBOX3 | P16989 | -1.554 | 0.0331 |
| 86 | A0A075B7D0 | A0A075B7D0 | -2.402 | 0.0004 |
| 87 | THEM4 | Q5T1C6 | -2.612 | 0.0305 |
| 88 | DTD1 | Q8TEA8 | -10.265 | 0.0115 |

**Note:** The table shows the protein cargo based on Log2 (Ratio) going from highest to lowest values.

**Supplementary Table 3.** Antibodies and reagents.

| Reagent | Company | Catalog # | Dilution |
| --- | --- | --- | --- |
| **Antibody** |  |  |  |
| Alix | Cell Signaling Technology | 2171 | 1:1000 |
| CD9 | Biolegend | 312102 | 1:1000 |
| Flotillin 1 | abcam | Ab133497 | 1:1000 |
| p-PHH3-antibody | Millipore Sigma | 06-570 | 1:500 |
| p-STAT1 (Ser727) | Cell Signaling Technology | 9177 | 1:200 |
| Guinea pig anti-Insulin | DAKO | A0564 | 1:1000 |
| Guinea pig anti-Insulin | abcam | Ab195956 | 1:1000 |
| Mouse anti-Glucagon | abcam | Ab10988 | 1:500 |
| Alexa Fluor 488 Goat α-Rabbit | Life Technologies | A11034 | 1:250 |
| Alexa Fluor 594 Goat α-Guinea Pig | Life Technologies | A11076 | 1:250 |
| Alexa Fluor 594 Goat α-Rabbit | Life Technologies | A11037 | 1:250 |
| Alexa Fluor 488 Goat α-Guinea Pig | Life Technologies | A11073 | 1:250 |
| Alexa Fluor 488 Goat α-Mouse | Life Technologies | A11029 | 1:250 |
| HRP Donkey α-Guinea Pig | Jackson ImmunoResearch | 706-035-148 | 1:500 |
| Anti human - CD3 | Biolegend | 317326 | - |
| Anti human - CD4-PE | Biolegend | 300508 | - |
| Anti human - CD8-Brilliant Violet 785 | Biolegend | 301046 | - |
| Anti human - CD25-Brilliant Violet 421 | Biolegend | 302630 | - |
| Anti human – CD69-PE/Cyanin7 | Biolegend | 310912 | - |
| **Reagents** |  |  |  |
| TNF-α | R&D Systems | 210-TA-020/CF | 1000 IU/ml |
| INF-γ | R&D Systems | 285-IF-100/CF | 1000 IU/ml |
| IL-1β | R&D Systems | 201-LB-005/CF | 100 IU/ml |
| CFSE | Thermo Fisher Scientific | C1157 | NA |
| Human Trustain FcX | Biolegend | 422302 | NA |
| Emapalumab | Selleckchem | A2041 | NA |
| Dynasore | Selleckchem | S8047 | NA |
| GW4869 | Cayman Chemical Company | 13127 | NA |
| Sirolimus | Selleckchem | S1039 | NA |
| Tacrolimus | Selleckchem | S5003 | NA |
| Collagenase P | Roche | 11249002001 | NA |
| Histopaque | Millipore Sigma | 10771 | NA |
| DeadEnd Fluorimetric TUNEL Kit | Promega | G3250 | NA |
| IFN-γ ELISA kit | Invitrogen | KAC1231 | NA |
| DAB Peroxidase Substrate Kit | Victor Labs | SK-4100 | NA |
| PKH26, Cell membrane label | Millipore Sigma | PKH26L | NA |
| DAPI (4',6-Diamidino-2-Phenylindole, Dilactate) | Life Technologies | D3571 | 1:100 |
| Mouse insulin kit | Mercodia | 10-1249-01 | NA |
| ExoQuick™, Exosome Precipitation Solution | System Biosciences | EXOQ5A-1 | NA |
| Size Exclusion Chromatography Column, 70 nm | IZON | ICO70 | NA |
| Deuterium Oxide (D<sub>2</sub>O) | Millipore Sigma | 151882 | NA |
| Sucrose | Millipore Sigma | G7021 | NA |
| Thromboplastin D | Fisher Scientific | 17-606-5 | NA |
| Hematoxylin | Sigma-Aldrich | HHS16-500 | NA |
| Eosin | Sigma-Aldrich | RO3040 | 5% |
| Hydrogen Peroxide | Sigma-Aldrich | H1009 | 1:10 |
| Triethylammonium bicarbonate (TEAB) | Thermo Fisher Scientific | PI90114PM | NA |
| (2-Carboxylethyl) phosphine (TCEP) | Thermo Fisher Scientific | T2556 | NA |
| Iodoacetamide | Thermo Fisher Scientific | AC122275000 | NA |
| Trypsin/LysC | Promega | V5071 | NA |

**Supplementary Table 4.** Characteristics of cadaveric human islet donors.

| Unique Identifier | Donor Age (Y) | Donor Sex (M/F) | Donor BMI (Kg/m<sup>2</sup>) | Donor HbA1c (%) | Source of Islets | Islet Isolation Center | Donor History of Diabetes |
| --- | --- | --- | --- | --- | --- | --- | --- |
| SAMN13710155 | 18 | M | 36.9 | 5.1 | IIDP | Southern California Islet Cell Resource Center, City of Hope | No |
| SAMN41299360 | 20 | M | 24.0 | 5.1 | IIDP | University of Arizona | No |
| SAMN44269935 | 29 | M | 26.5 | 5.4 | IIDP | University of Wisconsin | No |
| SAMN08768704 | 37 | F | 31.1 | NA | IIDP | University of Miami | No |
| SAMN41218531 | 38 | M | 29.1 | 5.2 | IIDP | Prodo Labs | No |
| SAMN23958504 | 42 | M | 37.9 | 5.7 | IIDP | Prodo Labs | No |
| SAMN37350251 | 43 | F | 29.9 | 5.2 | IIDP | University of Wisconsin | No |
| SAMN25860453 | 44 | M | 26.8 | 6.0 | IIDP | University of Pennsylvania Islet Transplant Center | No |
| SAMN08799544 | 45 | F | 32.1 | NA | IIDP | University of Miami | No |
| SAMN23481790 | 48 | M | 37.4 | 5.7 | IIDP | Loya Medicine, IL | No |
| SAMN08971735 | 50 | M | 32.3 | NA | IIDP | Prodo Labs | No |
| SAMN08612554 | 51 | F | 32 | NA | IIDP | University of Miami | No |
| SAMN36704819 | 52 | M | 25.9 | 4.9 | IIDP | Imagine Islet Center, Imagine Pharma | No |
| SAMN37871873 | 59 | M | 41.7 | 5.4 | IIDP | VCU Islet Cell Processing Lab | No |
| SAMN08768698 | 60 | M | 26.6 | NA | IIDP | Prodo Labs | No |
| Hu1149 | 18 | M | 24.4 | 4.8 | City of Hope | Southern California Islet Cell Resource Center, City of Hope | No |
| Hu1232 | 45 | F | 25.2 | 4.9 | City of Hope | Southern California Islet Cell Resource Center, City of Hope | No |
| Hu1226 | 47 | F | 31.6 | 5.5 | City of Hope | Southern California Islet Cell Resource Center, City of Hope | No |
| Hu1228 | 50 | F | 30.5 | 4.9 | City of Hope | Southern California Islet Cell Resource Center, City of Hope | No |
| Hu1250 | 50 | M | 36.3 | 4.9 | City of Hope | Southern California Islet Cell Resource Center, City of Hope | No |
| Hu1231 | 54 | F | 26.8 | 5.7 | City of Hope | Southern California Islet Cell Resource Center, City of Hope | No |
| Hu1289 | 54 | M | 22.3 | 5.3 | City of Hope | Southern California Islet Cell Resource Center, City of Hope | No |
| Hu1220 | 57 | M | 22.6 | 8.3 | City of Hope | Southern California<br>Islet Cell Resource<br>Center, City of Hope | No |
| Hu1301 | 60 | M | 27.8 | 5.6 | City of Hope | Southern California<br>Islet Cell Resource<br>Center, City of Hope | No |
| Hu1222 | 66 | M | 29.9 | 5.8 | City of Hope | Southern California<br>Islet Cell Resource<br>Center, City of Hope | No |
| HP-17321 | 25 | M | 25.8 | 5.7 | Prodo Labs | Prodo Labs | No |
| HP-23270 | 34 | M | 27.01 | 5.5 | Prodo Labs | Prodo Labs | No |
| HP-23258 | 39 | M | 26.89 | 5.2 | Prodo Labs | Prodo Labs | No |
| HP-24152 | 47 | F | 25.51 | 4.9 | Prodo Labs | Prodo Labs | No |

## Notes

### Competing Interest Statement

The authors have declared no competing interest.

